# Functional framework of the kinetochore and spindle assembly checkpoint in Arabidopsis thaliana

**DOI:** 10.1101/2024.11.04.621965

**Authors:** Aladár Pettkó-Szandtner, Zoltán Magyar, Shinichiro Komaki

## Abstract

The kinetochore, critical for accurate chromosome segregation and genome stability in eukaryotes, comprises the Constitutive Centromere Associated Network (CCAN) and the KMN network. In animals, the CCAN associates with centromeric nucleosomes throughout the cell cycle, while the KMN network assembles at kinetochores during M phase, binding spindle microtubules and serving as a platform for the spindle assembly checkpoint (SAC) complex. Despite conserved functions, kinetochore components vary across organisms. This study investigates the subcellular localization and interaction maps of core kinetochore components in *Arabidopsis thaliana*, finding that only CENP-C of the four conserved CCAN components localizes to kinetochores, while all KMN components consistently localize to the kinetochore throughout the cell cycle. Immunoprecipitation assays revealed interactions between core kinetochore proteins and regulators involved in DNA replication, histone modification, and chromatin remodeling, suggesting that the kinetochore may also function outside of M phase. Examining interactions between kinetochore and SAC components elucidates plant-specific SAC localization mechanisms providing a functional framework for understanding plant kinetochores and offering new insights into SAC regulation in plants.

## INTRODUCTION

The kinetochore is a large protein complex essential for accurate chromosome segregation during cell division. The M phase checkpoint consisting of the Spindle Assembly Checkpoint (SAC) and the Chromosomal Passenger Complex (CPC) (Lara-Gonzalez et al., 2012) monitors this process. The kinetochore, formed at the chromosome’s centromere, comprises three layers: the inner centromere, the inner kinetochore, and the outer kinetochore.

The inner centromere is a specialized chromatin region between sister centromeres, where the CPC monitors interkinetochore tension and prevents chromosome missegregation (Carmena et al., 2012). The proteins that localize to the inner kinetochore throughout the cell cycle are known as the Constitutive Centromere-Associated Network (CCAN). In vertebrates, the CCAN consist of 16 proteins organized into five groups: CENP-C, CENP-H/I/K/M, CENP-L/N, CENP-O/P/Q/R/U, and CENP-S/T/W/X. CENP-C and CENP-N directly interact with CENP-A, a centromere-specific histone H3 variant, while CENP-T binds to centromere DNA. These interactions are essential for assembling the CCAN at the inner kinetochore (McAinsh and Meraldi, 2011).

The outer kinetochore, extending from the inner kinetochore directly binds spindle microtubules. It consist of 10 core proteins forming three subcomplexes, collectively known as the KMN network: KNL1/ZWINT1 (KNL1 complex: KNL1C), MIS12/NNF1/NSL1/DSN1 (MIS12 complex: MIS12C), and NDC80/NUF2/SPC24/SPC25 (NDC80 complex: NDC80C) (Varma and Salmon, 2012). These subcomplexes facilitate spindle microtubule attachment and are crucial for force generation and signaling during chromosome movement.

The KMN network is recruited to the kinetochore via two independent pathways, mediated by CENP-C and CENP-T, in a sequential process throughout the cell cycle (Gascoigne et al., 2011). Initially, MIS12C and KNL1 are recruited to kinetochores during S phase (Gascoigne and Cheeseman, 2013). NDC80 then localizes to the kinetochore following nuclear envelope breakdown (NEB) in the M phase. Mis12C directly binds to CENP-C (Screpanti et al., 2011), creating a binding site for NDC80 aiding in KMN network assembly. In parallel, CENP-T also directly binds to NDC80C, and facilitating the assembly of the KMN network (Nishino et al., 2013) (Huisin’T Veld et al., 2016a).

Although kinetochore functions are critical for chromosome segregation, a fundamental eukaryotic process, its composition varies significantly among organisms (Hooff et al., 2017). In *Caenorhabditis elegans* (*C. elegans*) and *Drosophila melanogaster* (*D. melanogaster*), most CCAN components, including CENP-T, are absent. Thus, CENP-C alone mediates the CCAN and the KMN network interactions in these model organisms. Furthermore, many holocentric insects, early diverging fungi, and kinetoplastids lack CENP-A (Ishii and Akiyoshi, 2022). Therefore, obtaining kinetochore information from more species is crucial to understand evolutionary changes in kinetochore factors and formation.

The SAC is a surveillance system that ensures equal chromosome segregation during cell division. Core SAC components, including BUBR1, BUB3, MAD1, MAD2, and the kinases BUB1 and MPS1, are conserved from yeast to animals (London and Biggins, 2014a). Evolution has led to specialization and domain reshuffling among these core components such as BUB1-type proteins (Tromer et al., 2016). In a previous study, we found that Arabidopsis’s BUB1-type paralogs, BMF1, BMF2, and BMF3, show domain reorganization and exhibit functional differences from animal BUB1 (Komaki and Schnittger, 2017). Additionally, Arabidopsis BUB3.3 is more involved in chromosome congression than in the plant SAC (Lampou et al., 2023). Thus, despite SAC component conservation, SAC function and regulation likely vary among organisms.

The SAC complex is assembled on kinetochores unattached to microtubules. In yeast and humans, MPS1 kinase binds to NDC80,phosphorylates the MELT motifs in KNL1, and initiates SAC complex assembly at kinetochores (Shepperd et al., 2012)(Yamagishi et al., 2012)(London et al., 2012). The phosphorylated KNL1 facilitates the formation of the mitotic checkpoint complex (MCC), inhibiting the anaphase promoting complex/cyclosome (APC/C) to prevent premature chromosome segregation.

Sequence analyses indicate that plant genomes have lost many CCAN components, but retain KMN components (Yamada and Goshima, 2017)(Hooff et al., 2017). In the moss *Physcomitrella patens* (*P. patens*), only CENP-C (PpCENP-C) localizes to the kinetochore among conserved CCAN components, but it is absent post M phase, unlike in other organisms. Moss KMN components localize to kinetochores during M phase, similar to yeast and animals (Kozgunova et al., 2019). While some functions and localizations of kinetochore components in flowering plants are known (Girard et al., 2014)(Shin et al., 2018)(Li et al., 2021)(Allipra et al., 2022), comprehensive analyses are lacking , leaving plant kinetochore details unclear. Previously, we showed that the SAC in Arabidopsis is rapidly deactivated during severe stress, impacting ploidy levels and plant evolution (Komaki and Schnittger, 2017). However, the exact localization and regulation of the plant SAC at the kinetochore remained unknown.

This study provides a comprehensive localization and interaction map of core kinetochore components in Arabidopsis, detailing their assembly and functions beyond the M phase. We also identified unique plant-specific interactions between SAC and kinetochore factors, offering insight into the kinetochore structure and SAC regulation mechanism in plants.

## RESULTS

### Spatiotemporal localization of core kinetochore components in Arabidopsis

Genomic analysis identified four CCAN components and ten KMN network components in *A. thaliana* (Table 1). Each component is conserved as a single gene, except for NLS1 (NSL1.1 and NSL1.2), SPC24 (SPC24.1 and SPC24.2) and ZWINT1 (ZWINT1.1 and ZWINT1.2). To investigate the spatiotemporal localization of these core kinetochore proteins, at least one gene family member was fused with the green fluorescent protein (GFP) coding sequence and transformed into Arabidopsis plants. The resulting lines revealed the localization patterns of these CCAN and KMN components in the root meristem (Fig. 1; Supplemental Movies S1-S15).

**Table 1.**
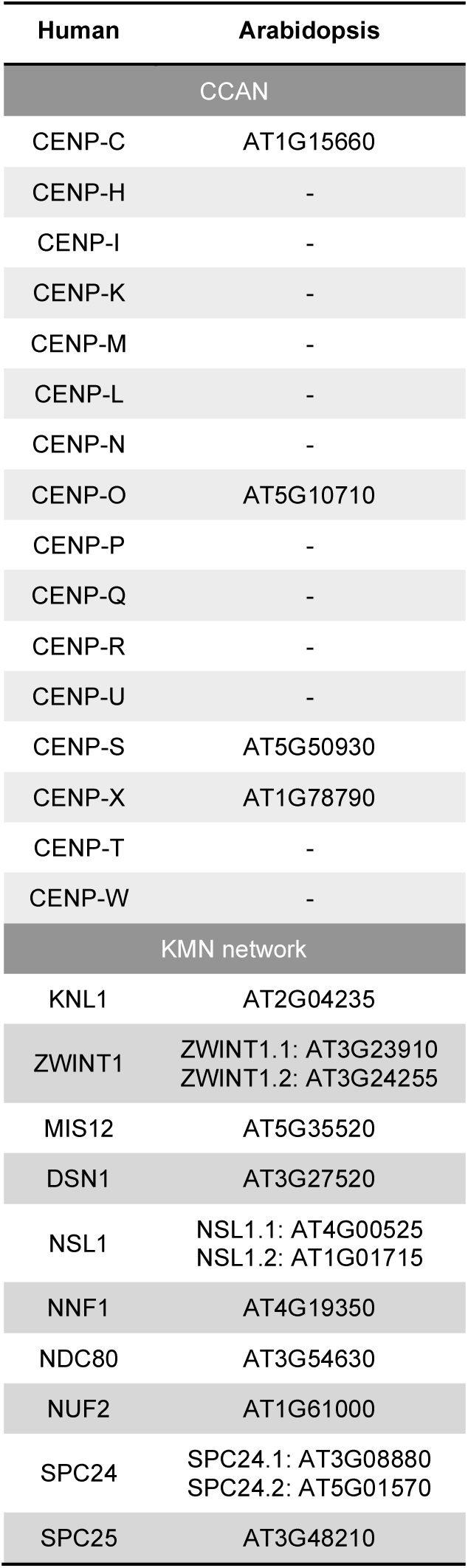
A comparative gene list of core kinetochore components in human and Arabidopsis.

**Fig 1.**
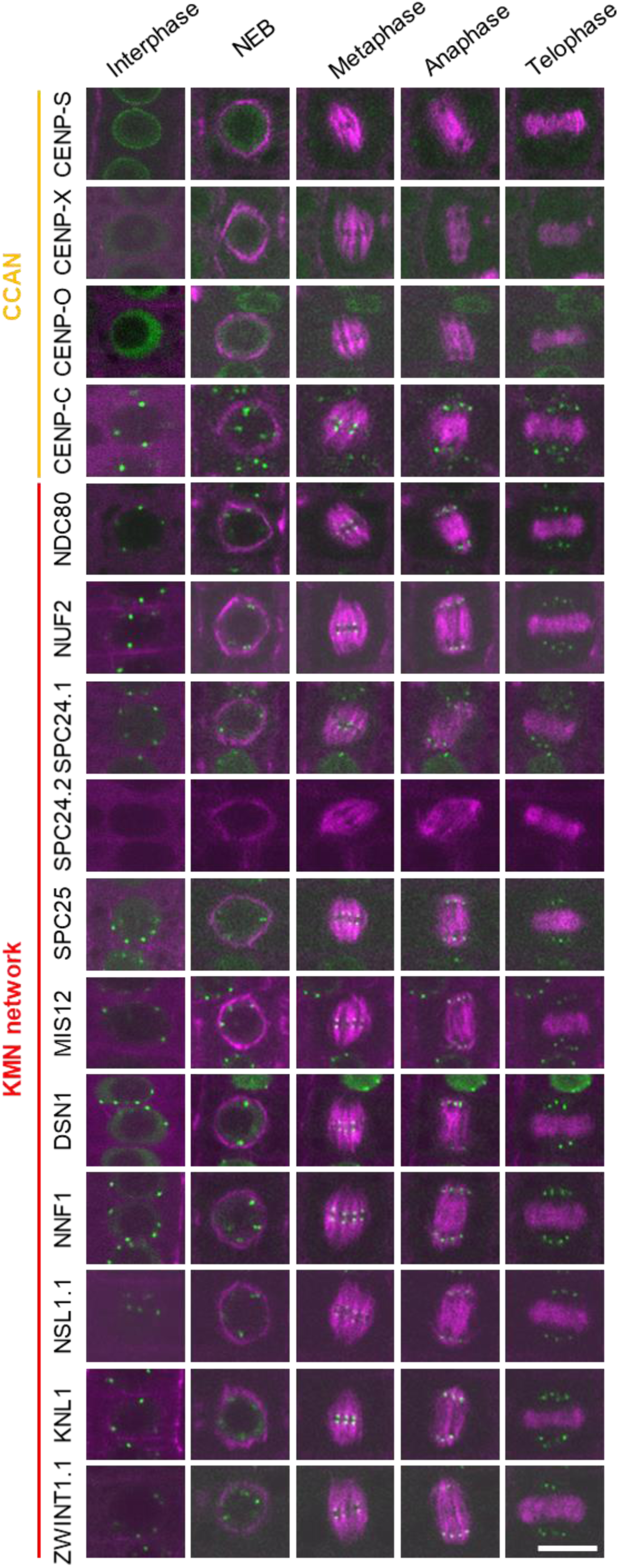
Subcellular localization of kinetochore component proteins during the cell cycle. Each kinetochore reporter line (green) was crossed with TagRFP:TUA5 (magenta) expressing plants to visualize microtubule structures. For live imaging, root tips of five-day-old seedlings were used. Scale bar, 10 µm.

The conserved CCAN component CENP-C localizes to Arabidopsis kinetochores throughout the cell cycle, consistent with findings in *P. patens*, while other CCAN components only exhibit kinetochore localization in animals (Kozgunova et al., 2019)(Fig. 1; Supplemental Movies S1-S4). CENP-S localized to the nuclear envelope during interphase and to the cytoplasm during mitosis (Fig. 1; Supplemental Movie S1). Additionally, CENP-X and CENP-O remained in the cytoplasm throughout the cell cycle (Fig. 1; Supplemental Movies S2-S3).

The KMN network, composed of KNL1C, MIS12C, and NDC80C, localizes to the outer kinetochore. In animal cells, the mature KMN network is present at kinetochores only during mitosis. Although SPC24.2 was not detected in root cells, other KMN components localized to kinetochores during both interphase and mitosis (Fig. 1; Supplemental Movies S5-S15).

In summary, only CENP-C exhibited the typical localization pattern of CCAN components in Arabidopsis. Furthermore, all KMN components in Arabidopsis are consistently localized to the kinetochore, unlike in animals.

### Building the Arabidopsis core kinetochore interactome

To elucidate the interaction network of core kinetochore components in Arabidopsis, we conducted immunoprecipitation (IP) experiments using an antibody against GFP on all kinetochore-GFP lines, except SPC24.2:GFP , followed by mass spectrometry.

Each bait was purified at least twice from extracts of one-week-old seedlings. We identified bait-specific interactions by selecting proteins with at least two unique peptides detected in at least two independent experiments (see Materials and Methods for details). Potential background proteins were excluded based on their occurrence in the 10 negative controls, resulting in 863 unique proteins as potential interactors of the 14 Arabidopsis core kinetochore components (Fig. 2, A and C; Supplemental Table. S1). GO enrichment analysis revealed that the IP lists were over 10-fold enriched in proteins linked to ‘kinetochore organization’ (31.98-fold; p = 2.57E-03) and ’centromere complex assembly’ (15.99-fold; p = 1.77E-02) (Supplemental Table. S2).

**Figure 2.**
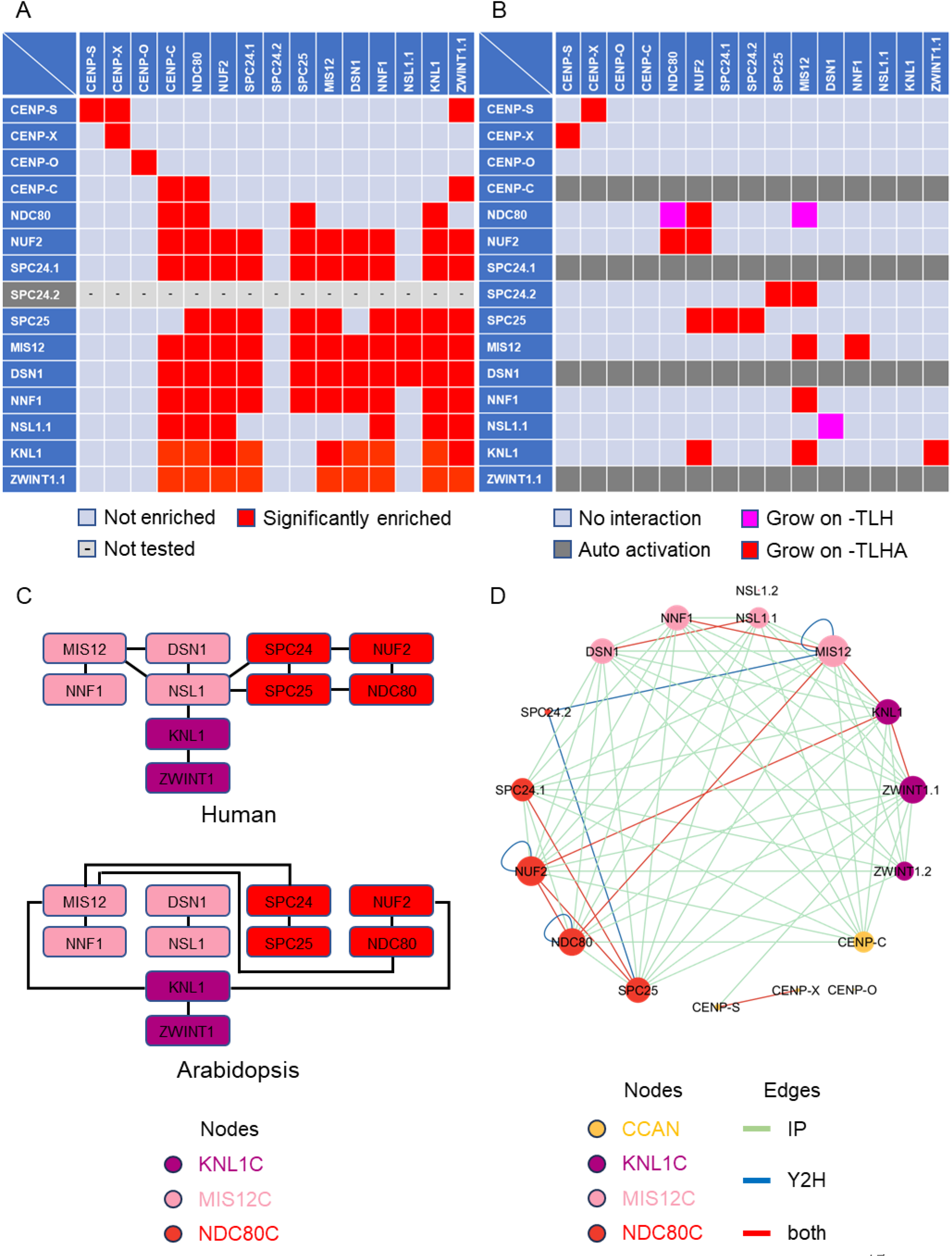
Interaction map of Arabidopsis core kinetochore components. A, Interaction among the core kinetochore components as revealed by IP. SPC24.2:GFP was not utilized in IP experiments, as no expression was observed in seedings. The corresponding IP data is available as Supplemental Table S1. B, Results of Y2H assays among core kinetochore components. Each yeast strain was spotted on SD plates without Trp and Leu (-TL; control media), without Trp, Leu, and His (-TLH; medium selection media), or without Trp, Leu, His, and Ade (-TLHA; severe selection media), and photographed after incubation at 30°C for three days. The interactions were classified according to yeast growth on different selection media in three categories. The color coding is as follows: pink indicates growth on -TLH; red indicates growth on -TLHA; light blue indicates growth rate not stronger than with mEGFP control; and dark grey indicates strong auto-activation observed. AD, GAL4-activation domain; BD, GAL4-DNA binding domain. The corresponding Y2H data is available as Supplemental Fig. S1. C, Comparison of binding patterns between KMN network factors in human and Arabidopsis. The binding model for Arabidopsis is based on Y2H results but it does not account for homodimerization. D, Cytoscape representation of the observed interaction network. The color and size of a node represent the subcomplex to which it belongs, and the rate of occurrence of each protein, respectively. The edge colors indicate the type of assay that detected the interaction: IP (green), Y2H (blue), and both methods (red).

To identify direct interactions among the IP results, we conducted a Y2H assay, testing 256 binary interactions among core kinetochore components in Arabidopsis. The cDNA of all kinetochore components and GFP, as a negative control, were cloned in pGBT9 (bait) and pGAD424 (prey) vectors and co-transformed in yeast cells with all 256 combinations (Fig. 2B and C; Supplemental Fig. S1). Since SPC24.2 cDNA was successfully amplified from flowering tissue, it was included it in the Y2H assay. Interactions were assessed by measuring growth on control (-TL), moderately selective (-TLH), and severely selective (-TLHA) media. However, CENP-C, SPC24.1, DSN1, and ZWINT1.1 with DB domain-containing constructs exhibited self-activation complicating the evaluation of their interactions.

### A comprehensive interaction map of the core kinetochore

We first examined the core kinetochore proteins. In animals, the CCAN is constantly localized to kinetochores and is essential for the formation of the KMN network (Hara and Fukagawa, 2018). In Arabidopsis, only CENP-C among the CCAN components localized to kinetochores (Fig. 1), co-purifying with many KMN components. However, other CCAN components like CENP-S, CENP-X, and CENP-O, were rarely detected in our IP samples (Fig. 2A and D). Only the CENP-S and CENP-X interaction was observed by both IP and Y2H (Fig. 2A, B and D).

Conversely, KMN network components were frequently detected in each other’s IP samples indicating a robust complex (Fig. 2A and D). Y2H results showed that the KNL1C, MIS12C, and NDC80C subcomplexes directly bound to each other through at least one component (Fig. 2B and C). However, the bridging components differed from animals, where NDC80C interacts with NSL1 of the MIS12C via SPC24-SPC25 (Petrovic et al., 2010) (Fig. 2B and C). In Arabidopsis, NDC80C interacts with

MIS12 via NDC80 and SPC24 (Fig. 2B and C). In animals, the KNL1C localizes to the kinetochore through KNL1 and MIS12’s NSL1 (Petrovic et al., 2010). In Arabidopsis, KNL1 binds to ZWINT1.1, MIS12 and NUF2 indicating localization via both MIS12C and NDC80C subcomplexes (Fig. 2B and C). Similar to animals, Arabidopsis MIS12C components formed MIS12-NNF1 and NSL1.1-DSN1 heterodimers (Fig. 2B and C). Additionally, Y2H results showed homodimerization of NDC80, NUF2, and MIS12 (Fig. 2B). Collectively, subcellular localization results indicate that the Arabidopsis KMN network form a complex at the kinetochore throughout the cell cycle, exhibiting distinct binding modes compared to animals.

### Identification of novel interactors with core kinetochore components in Arabidopsis

Our IP data set revealed numerous proteins previously unidentified as kinetochore interactors in plants. Focusing on cell cycle and/or cytoskeleton-related proteins under specific GO IDs: cell cycle (GO:0007049), cell division (GO:0051301), cell population proliferation (GO:0008283), chromosome segregation (GO:0007059), DNA replication (GO:0006260), chromosome organization (GO:0051276), histone binding (GO:0042393), chromatin organization (GO:0006325), cytoskeleton organization (GO:0007010), cytoskeletal protein binding (GO:0008092). Among 1712 Arabidopsis proteins 98 related to the cell cycle and cytoskeleton were identified in the IP samples (Table 2).

**Table 2.**
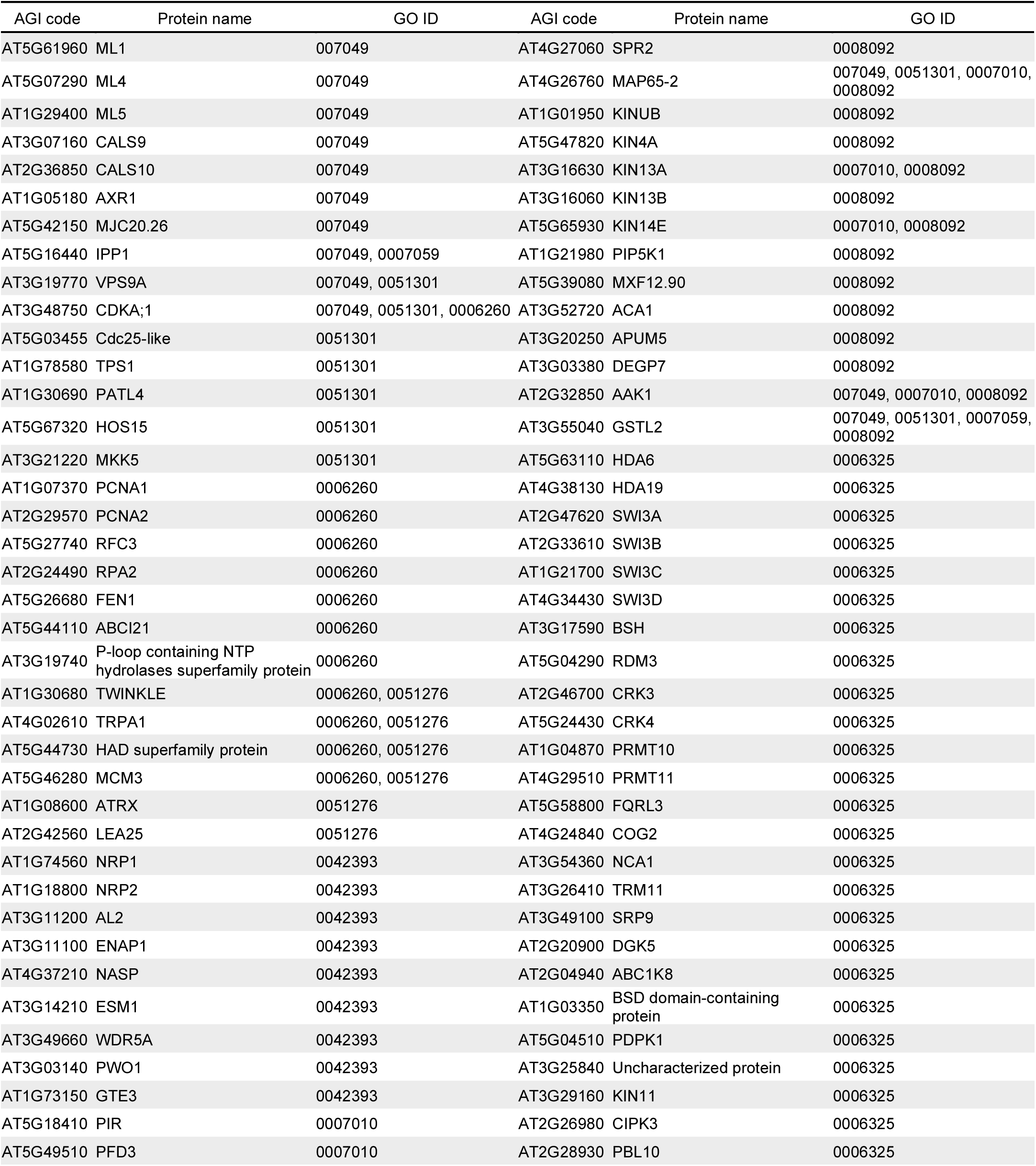
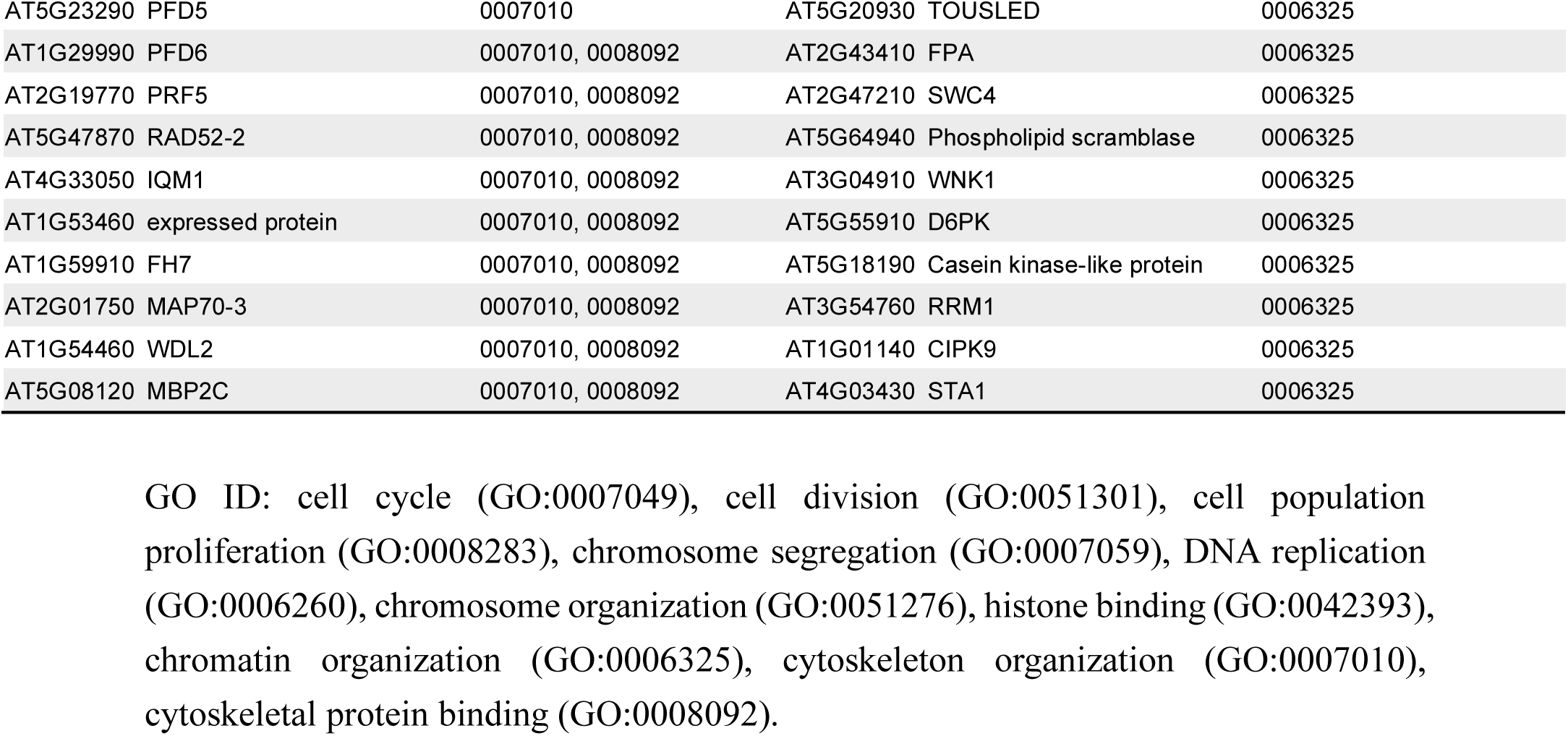
List of novel interactors with core kinetochore components in Arabidopsis.

CDKA;1, a key cyclin-dependent kinase (CDK) in Arabidopsis, copurified with core kinetochore proteins. KNL1, NUF2, and DSN1, were phosphorylated at [S/T]P CDK consensus sites (Supplemental Table S3), suggesting that CDKA;1 may be responsible for these modifications. The IP lists also include DNA replication factors (PCNA1, PCNA2, RFC3, RPA2, FEN1, and MCM3) and histone modification proteins (HDA6, HDA19, HOS15, and WDR5A), indicating a linkage between kinetochores and DNA replication. Additionally, subunits of the SWI/SNF complex (SWI3A, SWI3B, SWI3C, SWI3D, and BSH) were frequently observed. The SWI/SNF complexes, known for chromatin remodeling and transcription regulation, have also been localized to yeast kinetochores (Xue et al., 2000), suggesting a similar role in plants. Moreover, several kinesins (KINUB, KIN4A, KIN13A, KIN13B, and KIN14E) and microtubule-binding proteins (MBP2C, SPR2, WDL2, and MAP65-2) copurified with core kinetochore components, interacting with cortical microtubules and regulating cell morphology, thus emphasizing kinetochores’ broader functional significance beyond the M phase.

### SAC assembly at kinetochore

The SAC complex localizes to kinetochores, ensuring genome stability by facilitating proper chromosome segregation during mitosis and meiosis. In animals, kinase MPS1, a SAC component, localizes to kinetochores via NDC80, phosphorylates KNL1 and activates SAC signaling (London and Biggins, 2014a). SAC silencing involves the competition for NDC80 binding between spindle microtubules and MPS1 (Ji et al., 2015)(Hiruma et al., 2015). Therefore, the MPS1-NDC80 interaction is vital for both SAC activation and silencing. Our previous work showed that the Arabidopsis SAC components MPS1, BMF1, BMF3, MAD1, and MAD2 are present at kinetochores (Komaki and Schnittger, 2017). However, the specific kinetochore factors responsible for SAC component localization in Arabidopsis were unclear. We explored this by analyzing SAC and conserved kinetochore component interactions using Y2H assay (Fig. 3A; Supplemental Fig. S2). Surprisingly, Arabidopsis MPS1 binds directly to KNL1 not NDC80 (Fig. 3A), implying no competition between MPS1 and microtubules for NDC80 binding in Arabidopsis cells. In human cells, MPS1-phosphorylated KNL1 scaffolds BUB3-BUB1 and BUB3-BUBR1 complexes recruiting the MAD1-MAD2 complex to kinetochores through BUB1 kinase and MAD1 interaction mediated by a conserved domain 1 (CD1) in BUB1 (London and Biggins, 2014b). In Arabidopsis BUB1 and BUBR1 functions are divided among three proteins: BMF1 (BUB1 kinase domain), BMF2 (BUBR1-like), and BMF3 (BUB1 CD1-like domain)(Komaki and Schnittger, 2017). We demonstrated that BMF3 interacts with MAD1 via its CD1-like domain, and Arabidopsis MAD1 also binds core kinetochore components, including NDC80, NUF2, and KNL1 (Fig. 3A). Thus, the MAD1-MAD2 complex may localize to the kinetochore via BMF3-dependent and independent pathways.

**Fig 3.**
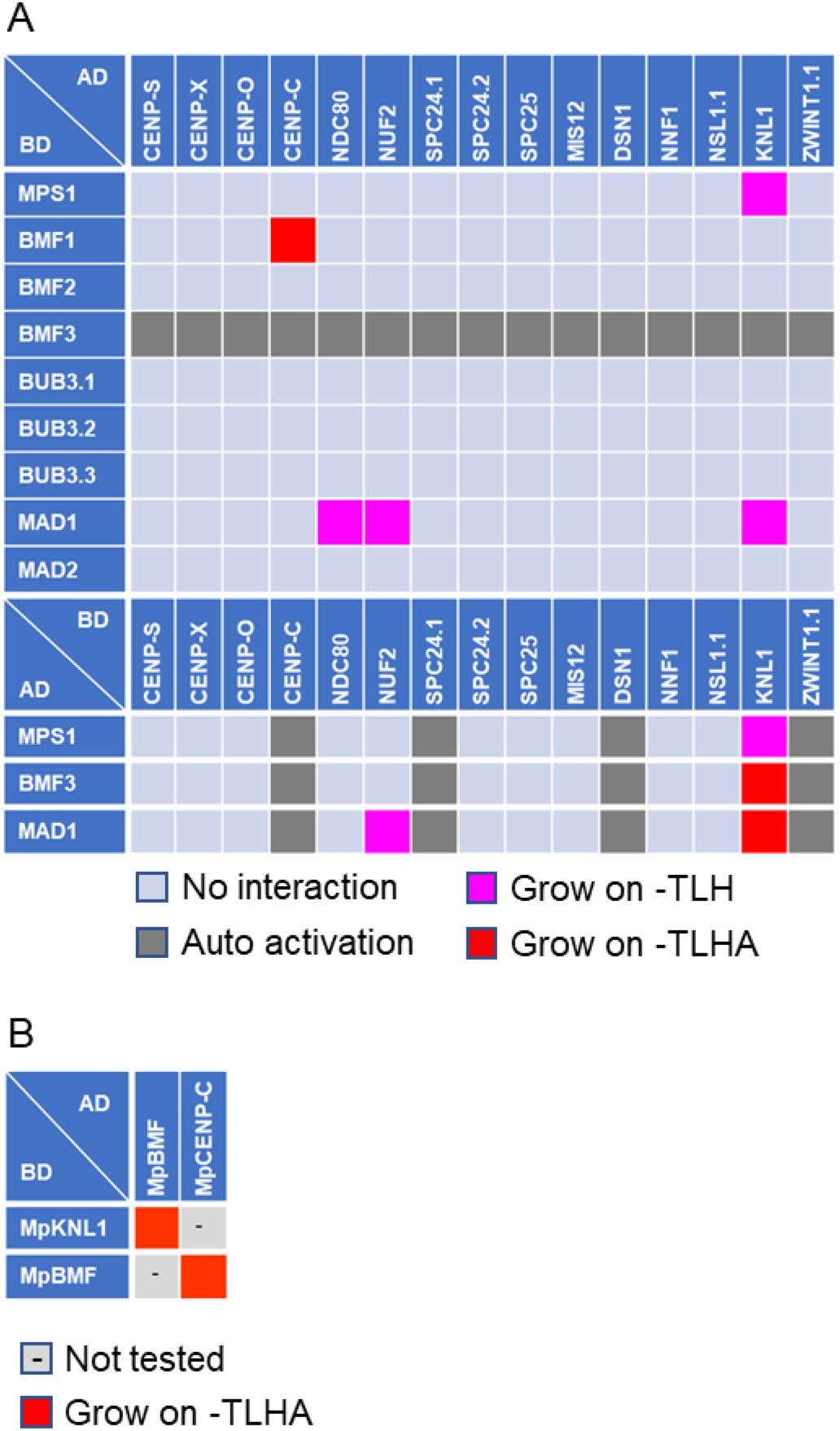
Interaction between core kinetochore and SAC components. A, Results of Y2H assays between Arabidopsis core kinetochore components and SAC components. B, Results of Y2H assays between *M. polymorpha* core kinetochore components (MpCENP-C and MpKNL1) and SAC component (MpBMF). Each yeast strain was spotted on SD plates without Trp and Leu (-TL; control media), without Trp, Leu, and His (-TLH; medium selection media), or without Trp, Leu, His, and Ade (- TLHA; severe selection media), and photographed after incubation at 30°C for three days. The interactions were classified according to yeast growth on different selection media in three categories. The color coding is as follows: pink indicates growth on -TLH; red indicates growth on -TLHA; light blue indicates growth rate not stronger than with mEGFP control; and dark grey indicates strong auto-activation observed. AD, GAL4- activation domain; BD, GAL4-DNA binding domain. The corresponding Y2H data is available as Supplemental Fig. S2.

To map SAC complex interactions, we investigated SAC component binding. Besides previously identified interactions (MAD1-BMF3, MAD1-MAD1, and MAD1-MAD2) (Komaki and Schnittger, 2017), we found that MPS1 and BMF2 form homodimers (Fig. 4; Supplemental Fig. S3). These findings provide a more detailed understanding of SAC assembly and kinetochore regulation.

**Fig 4.**
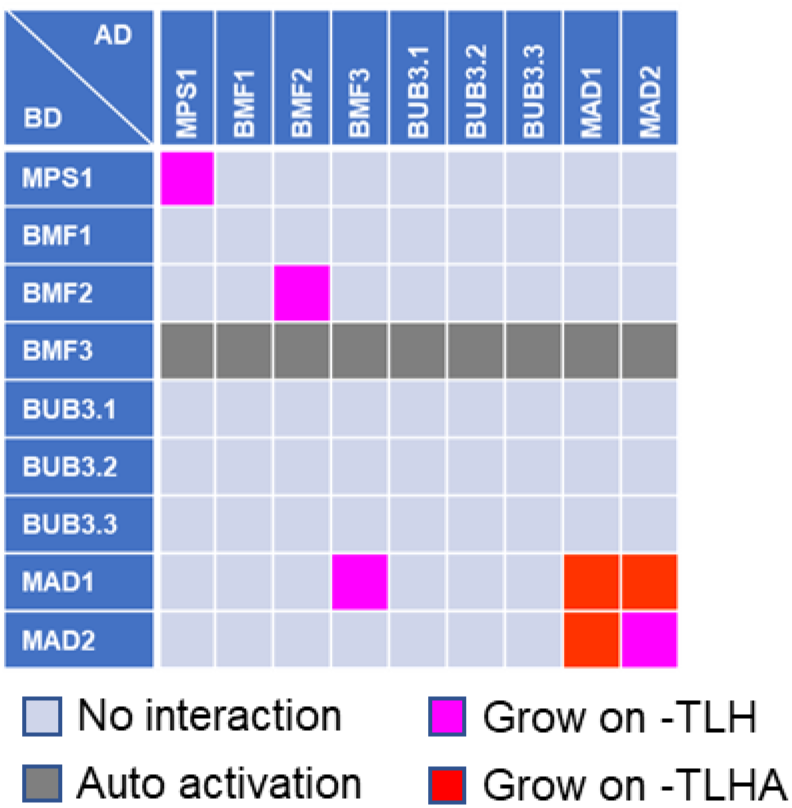
Interaction among Arabidopsis SAC components. Results of Y2H assays among Arabidopsis SAC components. Each yeast strain was spotted on SD plates without Trp and Leu (-TL; control media), without Trp, Leu, and His (-TLH; medium selection media), or without Trp, Leu, His, and Ade (-TLHA; severe selection media), and photographed after incubation at 30°C for three days. The interactions were classified according to yeast growth on different selection media in three categories. The color coding is as follows: pink indicates growth on -TLH; red indicates growth on -TLHA; light blue indicates growth rate not stronger than with mEGFP control; and dark grey indicates strong auto-activation observed. AD, GAL4- activation domain; BD, GAL4-DNA binding domain. The corresponding Y2H data is available as Supplemental Fig. S3.

### KNL1 function as an interaction hub between the kinetochore and the SAC

Our data show that three kinetochore components (MIS12, NUF2, and ZWINT1.1) and three SAC components (BMF3, MAD1, and MPS1) bind to KNL1 in plants (Fig. 2B and Fig. 3A). To identify KNL1’s interaction domain we examined truncated versions of the KNL1 construct using Y2H assay. Arabidopsis KNL1 has a eudicot-specific-domain (ESD), coiled-coil (CC), and RING-WD40-DEAD (RWD) domains. (Deng et al., 2024), BMF3 binds to the ESD containing region (Fig. 5; Supplemental Fig. S4), while, MAD1 and MPS1 bind to the ESD containing region and the region from amino acids 350 to 450, respectively (Fig. 5; Supplemental Fig. S4). Although this MPS1-binding region is not well conserved in KNL1 across plants, structural prediction using AlphaFold 2 and 3 suggest these regions form helices (Supplemental Fig. S5A and B)(Varadi et al., 2024). In animals, kinetochore components, ZWINT1 and NSL1, bind to the CC and the RWD domains, respectively (Kiyomitsu et al., 2011)(Petrovic et al., 2014). Our results demonstrated that Arabidopsis NUF2, MIS12, and ZWINT1.1 bind to the ESD containing region and the CC domain with ZWINT1.1 also binding to the RWD domain (Fig. 5; Supplemental Fig. S4).

**Fig 5.**
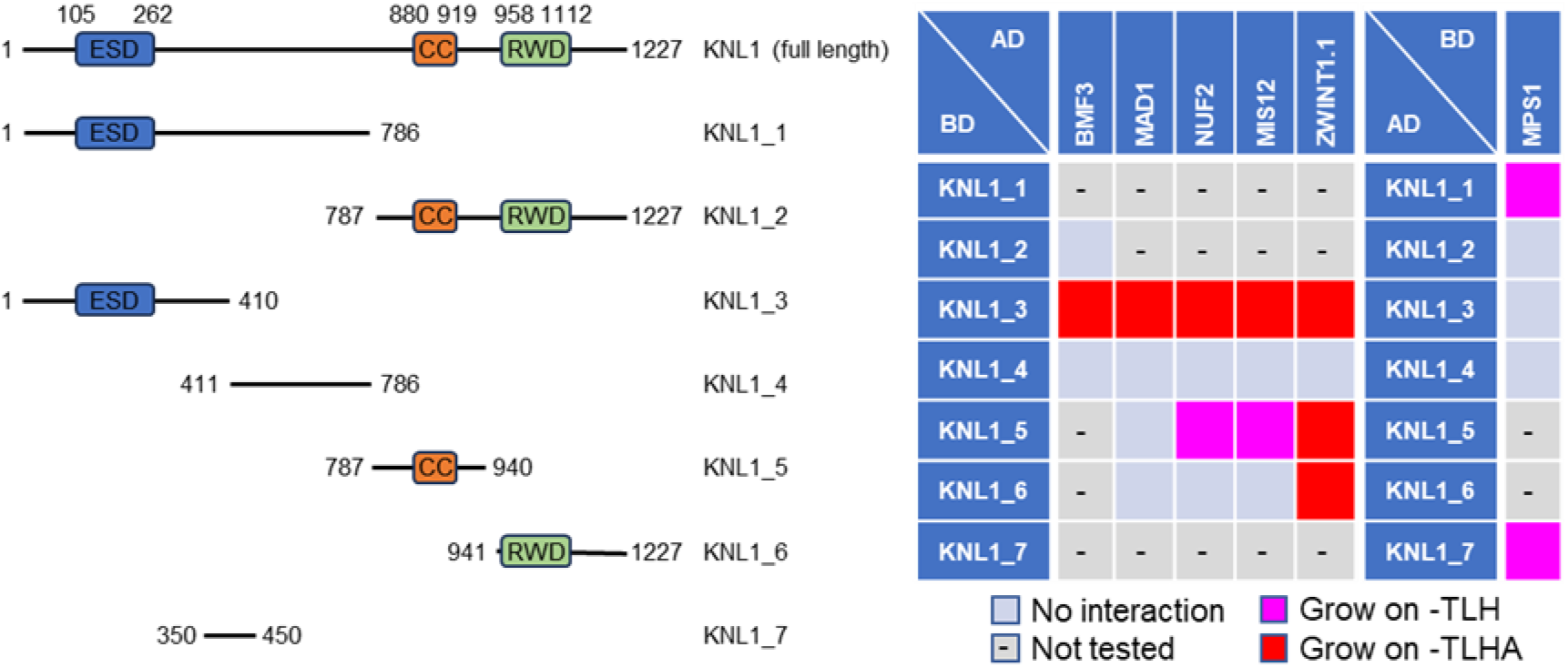
Identification of the interaction domain between KNL1 and its interactors. Y2H assays with the KNL1 fragments and its interactors. Each yeast strain was spotted on SD plates without Trp and Leu (-TL; control media), without Trp, Leu, and His (-TLH; medium selection media), or without Trp, Leu, His, and Ade (-TLHA; severe selection media), and photographed after incubation at 30°C for three days. The corresponding Y2H data is available as Supplemental Fig. S4.

These results indicate that KNL1 acts as a hub connecting kinetochores and the SAC through various pathways.

### BMFs loading system at the kinetochores

Su et al., (2021) have shown that maize KNL1 (ZmKNL1) binds to ZmBMF1 and ZmBMF2. In contrast, Arabidopsis KNL1 binds to BMF3, suggesting distinct BMFs loading system at the kinetochores in monocots and eudicots (Deng et al., 2024). Our Y2H results supports that Arabidopsis BMF3 binds to KNL1. Additionally, Arabidopsis BMF1, which has a BUB1 kinase domain, binds to CENP-C (Fig. 3A). Chlorophyta, such as *Micromonas pusilla* and *Volvox carteri*, possess a single BMF gene (Vleugel et al., 2012). To explore the evolutionary aspect of BMFs loading systems, we studied the moss *Marchantia polymorpha* (*M. polymorpha*), an early-diverging land plant model identifying one *BMF* family gene (*MpBMF*) that binds both MpCENP-C and MpKNL (Fig. 3B; Supplemental Fig. S2). This suggests the original plant BMF could bind to both proteins with functions splitting into BMF1 and BMF3 in angiosperms, each developing distinct binding partners.

## DISCUSSION

The kinetochore is essential for the equal segregation of chromosomes during cell division, a function vital for genome maintenance and conserved across eukaryotes from yeast to animals. However, its structure and components vary significantly among species. Although core kinetochore components in plants have been identified, the lack of molecular framework for these components has limited understanding. This study provides a comprehensive atlas of plant kinetochore factors along with their localization during the cell cycle (Fig. 6). Our findings suggest plant-specific kinetochore features significantly affect genome stability through SAC localization and broader functions beyond the M phase.

**Figure 6.**
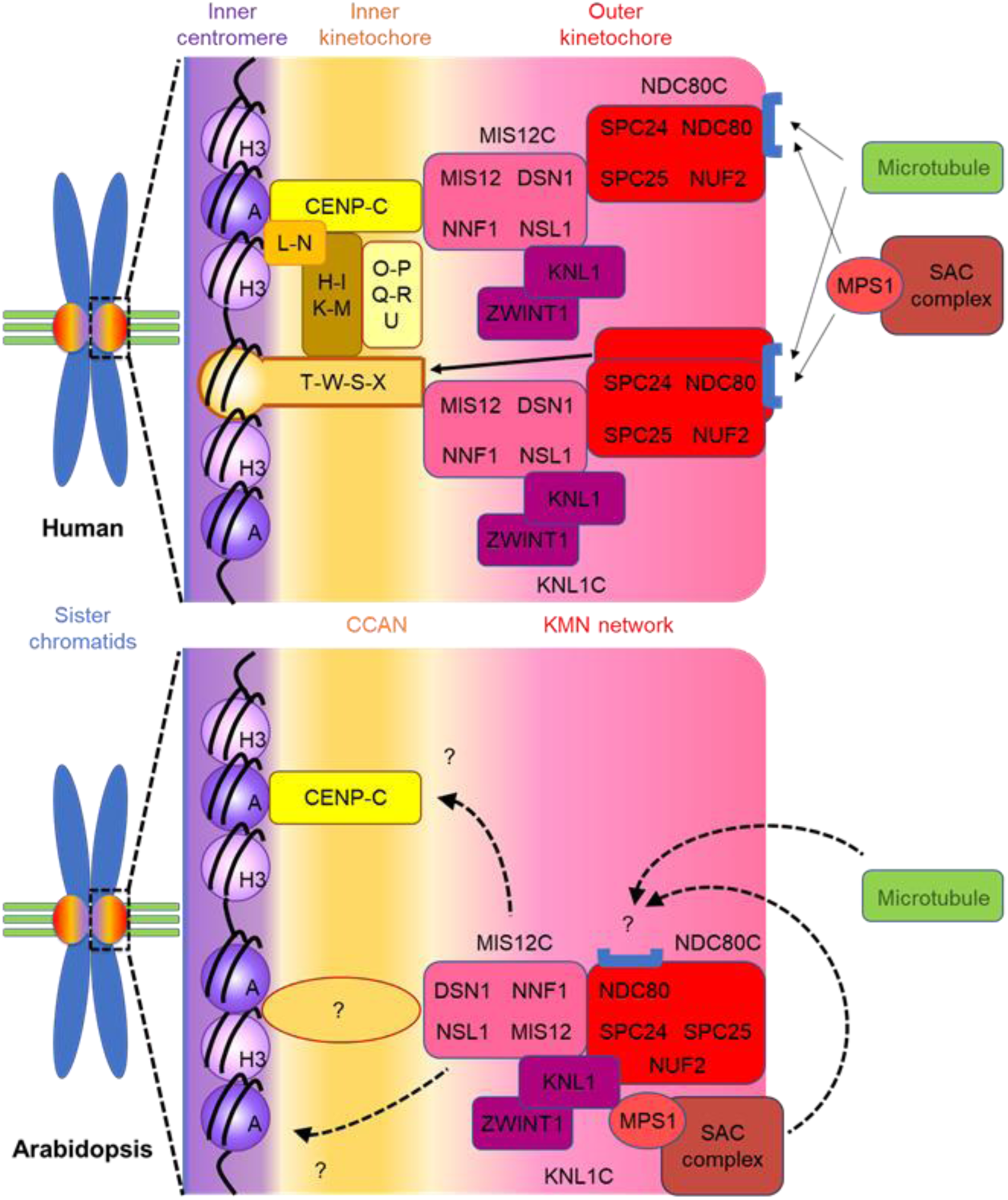
A schematic comparison of the interactions between core kinetochore and SAC components in human and Arabidopsis. The CCAN at the inner kinetochore is responsible for recruiting the KMN network through CENP-T and CENP-C in humans. Although the Arabidopsis KMN network is also recruited to kinetochores, the responsible protein(s) remain unknown. In humans, MPS1 and microtubules compete for the binding region of NDC80 at kinetochores. It is possible that this competition does not occur in Arabidopsis, given that Arabidopsis MPS1 binds KNL1 instead of NDC80.

Inner kinetochore proteins, known as CCAN, exhibit low sequence homology but high structural conservation between yeast and human (Yan et al., 2019). In both, CCAN components CENP-C and CENP-T bind directly to the centromere, essential for CCAN assembly into kinetochores. However, some organisms, such as *C. elegans* and *D. melanogaster* have lost most CCAN components, including CENP-T, relying solely on CENP-C for chromatin binding (Singleton, 2016). Arabidopsis similarly retains only CENP-C, CENP-S, CENP-X, and CENP-O, with only CENP-C localizing to the kinetochore and co-purifying with other components (Fig. 1 and Fig. 2, A-D). In *P. patens* moss, only CENP-C localizes to the kinetochore (Kozgunova et al., 2019). Thus, it is likely that chromosome-kinetochore binding via CENP-C is the primary pathway in plants, similar to *C. elegans* and *D. melanogaster*, with other CCANs have lost their kinetochore function in plant lineages. It is noteworthy that in human, CENP-S and CENP-X form heterodimers involved in DNA repair as well as kinetochore function. Our results show that Arabidopsis CENP-S and CENP-X, which form heterodimers do not localize to kinetochores, suggesting a role in DNA repair (Girard et al., 2014).

The CCAN components, CENP-C and CENP-T, independently recruit the KMN network to kinetochores in animals. The MIS12C directly binds the N-terminal region of CENP-C (1-71 aa in human) (Screpanti et al., 2011), recruiting NDC80 and the other KMN elements. Concurrently, CENP-T binds directly to NDC80C, aiding KMN network (Nishino et al., 2013)(Huisin’T Veld et al., 2016b)(Takenoshita et al., 2022). Although Arabidopsis CENP-C’s N-terminal amino acids show less conservation with humans (Supplemental Fig. S5), Arabidopsis CENP-C co-purifies with most KMN components, suggesting its role in KMN recruitment to the kinetochore. However, Y2H did not identify CENP-C’s direct interactor. This might be because both MIS12 and NNF1 are needed for CENP-C binding in some organisms (Richter et al., 2016)(Zhou et al., 2017) or MIS12C might bind directly to centromeres or involve plant-specific factors (Fig. 6). Consequently, further experiments with multiple proteins are necessary to identify CCAN and KMN interactions in Arabidopsis.

In animals, NDC80C’s kinetochore localization involves SPC24 and SPC25 binding to DSN1 and NSL1 of MIS12C, and KNL1C binding to NSL1 via KNL1. However, in Arabidopsis our Y2H results showNDC80C binds to MIS12 via NDC80 and SPC24. Furthermore, Arabidopsis KNL1 binds to MIS12 and NUF2 of NDC80C, indicating significant differences in KMN network construction compared to humans (Fig. 6). Protein alteration for kinetochore assembly is common; for example, DSN1 is absent in Drosophila but replaced functionally by Spc105R, a KNL1 homolog (Przewloka et al., 2009)(Liu et al., 2016). Further kinetochore plasticity is observed in protozoan parasites like apicomplexans and kinetoplastids, which lack many known kinetochore components and have highly divergent kinetochores (Akiyoshi and Gull, 2014)(Brusini et al., 2022). This diversity suggests that understanding of kinetochore evolution requires data from a broad range of organisms.

We found that the KMN network construction and kinetochore formation timing vary between animals and among plant lineages. In animals, MIS12C and KNL1 are recruited to the outer kinetochores during the S phase, with NDC80 released from the nucleus, restricting kinetochore formation to the M phase (Gascoigne and Cheeseman, 2013). In the moss *P. patens*, NDC80 localizes to the kinetochores only during the M phase (Kozgunova et al., 2019). Conversely, in Arabidopsis, all KMN components localize to the kinetochores throughout the cell cycle, affecting kinetochore functions and also the SAC localization potentially impacting SAC regulation (see further discussion below).

Our IP experiments indicate Arabidopsis core kinetochore factors may interact with previously unidentified non-canonical kinetochore proteins. CDKA;1, a key cell cycle regulatory kinase, was detected in most IP lists. We observed phosphorylation of KNL1, NUF2, and DSN1 at their CDK consensus sites, suggesting post-translational regulation by CDKA;1. In animals, CDK1 phosphorylates kinetochore proteins like CENP-C and NSL1, enhancing their binding to CENP-A and KNL1, respectively (Watanabe et al., 2019) (Lim-Manley et al., 2023). However, these specific amino acid residues are not conserved in plants, indicating different target kinetochore proteins.

Our IP data also identified proteins involved in DNA replication, including PCNA1, PCNA2, RFC3, RPA2, FEN1, and MCM3. PCNA1 and PCNA2 broadly co-purified with core kinetochore proteins, while RFC3, RPA2, FEN1, and MCM3 mainly co-purified with KMN network proteins. The continuous assembly of the Arabidopsis KMN network throughout the cell cycle, may serve as a scaffold for these DNA replication factors. Additionally, the IP list includes factors related to histone modification, chromatin remodeling, and microtubule-directed cell elongation, highlighting plant kinetochores’ significant roles throughout the cell cycle. Future research should verify the physiological significance of these interactions.

The SAC is crucial for genome stability, ensuring equal chromosome segregations during cell division. In animals, MPS1’s localization at the kinetochore, where it binds directly to NDC80, activates the SAC signal. We have previously shown that continuous stress inactivates the Arabidopsis SAC, leading to mitosis without nuclear division and resulting in polyploid cells (Komaki and Schnittger, 2017). This plant-specific SAC regulation may impact plant evolution by affecting ploidy levels. The exact SAC regulatory mechanisms in Arabidopsis are unclear, but MPS1 localizes to the kinetochore throughout the cell cycle, unlike in animals (Komaki and Schnittger, 2017). This study suggests that the Arabidopsis KMN network is consistently directed to the kinetochore, which may explain the persistent localization of MPS1. Therefore, the phosphorylation activity of MPS1, rather than its localization, is the primary SAC regulatory mechanism in plants. Interestingly, in the moss *P. patens* both MPS1 and the KMN network localize to the kinetochore only during the M phase (Kozgunova et al., 2019), indicating differences in SAC regulation between mosses and flowering plants. Furthermore, scaffold proteins for MPS1 at kinetochores differ between animals and Arabidopsis. In yeast and animals, MPS1 and microtubules compete for NDC80 binding at kinetochores and microtubule binding s to NDC80, displacing MPS1, is a known SAC silencing mechanism (Ji et al., not NDC80 (Fig. 3A and Fig. 5). In addition, MPS1 signals persist at anaphase (Komaki and Schnittger, 2017), suggesting a distinct SAC silencing pathway in plants.

Previous studies and our findings reveal that in monocots, KNL1 binds to BMF1 and BMF2, while in dicots, it binds to BMF3 (Fig. 3A) (Su et al., 2021)(Deng et al., 2024). In yeast and animals, KNL1’s conserved MELT motif , phosphorylated by MPS1, interacts with BUB1, analogous to BMF1 (Caldas and DeLuca, 2014). However, since plant KNL1 lacks a conserved MELT motif (Tromer et al., 2015), it is crucial to investigate if MPS1 activity is required for KNL1-BMFs binding. We also found Arabidopsis BMF1 binds to CENP-C. Remarkably, *M. polymorpha*’s sole BMF-like protein binds both KNL1 and CENP-C, hinting that the function of BMF may have split into multiple proteins during plant evolution. Although the last eukaryotic common ancestor (LECA) likely had a single BUB1 family protein (referred to as MadBub)(Suijkerbuijk et al., 2012), no MadBub-CENP-C interactions are reported. Thus, the BMF-CENP-C interaction might be plant specific. Further biological and bioinformatics studies are needed to verify this hypothesis.

In conclusion, our analysis highlights plant-specific interactions between kinetochore components and kinetochore-SAC components in Arabidopsis. We also discovered that core kinetochore components interact with factors functioning outside of M phase, suggesting that plant kinetochores might operate throughout the cell cycle. The universality of these characteristics across plants and their role in genome plasticity remain unclear due to limited studies on plant kinetochores. However, a recent report found key kinetochore and SAC components missing in *Cuscuta* genus parasitic plants, suggesting dynamic rearrangements in plant kinetochores (Neumann et al., 2023). Further research is needed to understand plant-specific SAC regulation and its potential impact on genome plasticity.

## MATERIALS AND METHODS

### Plant Materials and Growth Conditions

The *Arabidopsis thaliana* accession Columbia (Col-0) was used as the wildtype in this study. Plants were grown on a solid medium containing half-strength Murashige and Skoog (MS) salts, 1% (w/v) sucrose and 1.5% (w/v) agar in a growth chamber (16h day/8h night at *22°C)*.

### Plasmid Construction and Transgenic Plants

To create kinetochore markers, the genomic fragment of each kinetochore gene was amplified with *attB* sites by PCR and cloned into *pDONR221*. The *Sma*I site (*Nae*I site for *NNF1*) was inserted in front of the start codon (*CENP-C* and *NDC80*) or the stop codon (the other genes) of the constructs. The resulting construct was linearized by *Sma*I digestion and was ligated to the monomeric enhanced *GFP* (*mEGFP*) gene, followed by LR recombination reactions with the destination vector *pGWB501*(Nakagawa et al., 2007). Primer pairs for plasmid construction are listed in Supplemental Table S4. Transgenic plants were generated by the floral dip method (Clough and Bent, 1998). The constructs were introduced into *Agrobacterium tumefaciens* strain GV3101 (pMP90) and used for plant transformation.

### Immunoprecipitation

150-200 transgenic seedlings were harvested, frozen in liquid nitrogen and ground with TissueLyser (Qiagene) (50Hz, 4x 30 sec). Total proteins from seedlings were extracted as described (Henriques et al., 2010). Total protein extracts (2-4mg/IP) were immunopurified using anti-GFP antibody coupled to 50 nm size magnetic beads (MACS® Technology, Miltenyi) with a method modified from (Hubner et al., 2010) (Horvath et al., 2017), reduced with 10 mM DTT alkylated with 22 mM IAM digested in column with trypsin (Promega), and analyzed in a single run on the mass spectrometer.

### Mass spectrometry

The digestion mixtures were acidified and transferred to a single-use trapping mini­column (Evotip; 1/8 of the samples) and then analyzed with a data-dependent LC-MS/MS method using an Evosep One (LC: 15 SPD; MS1: R:120,000) on-line coupled to a linear ion trap-Orbitrap (Orbitrap-Fusion Lumos, Thermo Fisher Scientific) mass spectrometer operating in positive ion mode. Data acquisition was carried out in data-dependent fashion, multiply charged ions were selected in cycle-time from each MS survey scan for ion-trap HCD fragmentation (MS spectra were acquired in the Orbitrap (R=60000) while MSMS in the ion-trap).

### Data interpretation

Raw data were converted into peak lists using the in-house Proteome Discoverer (v 1.4) (Guan et al., 2011) and searched against the Swissprot database (downloaded 2019/6/12, 560292 proteins) using the Protein Prospector search engine (v5.15.1) with the following parameters: enzyme: trypsin with maximum 1 missed cleavage; mass accuracies: 5 ppm for precursor ions and 0.6 Da for fragment ions (both monoisotopic); fixed modification: carbamidomethylation of Cys residues; variable modifications: acetylation of protein N-termini; Met oxidation; cyclization of N-terminal Gln residues allowing maximum 2 variable modifications per peptide. Acceptance criteria: minimum scores: 22 and 15; maximum E values: 0.01 and 0.05 for protein and peptide identifications, respectively. Another database search was also performed using the same search and acceptance parameters except that Uniprot.random.concat database (downloaded 2022/07/20) was searched with *Arabidopsis thaliana* species restriction (136466 proteins) including additional proteins identified from the previous Swissprot search (protein score>50).

### Confocal microscopy

The root tips of five-day-old seedlings were utilized for live cell imaging. Samples were placed in glass-bottom dishes and covered with a solid medium containing half-strength MS salts, 1% sucrose, and 1.5% agar. Confocal images were acquired using a Leica TCS SP8 inverted confocal microscope with a HC PL APO 63x/1,20 CS2 water immersion objective. mEGFP and TagRFP-T were excited using 488 nm and 555 nm laser, respectively. Images were obtained at 20-second intervals and corrected for image drift by the StackReg plugin (Rigid Body option) for ImageJ version 1.54f.

### Yeast Two-Hybrid (Y2H) Assay

To create the Y2H constructs, the cDNA fragments of each gene were amplified by PCR and cloned into *pENTR4* by the SLiCE method or amplified with *attB* sites by PCR and cloned into *pDONR221*. The *MPS1*, *BMF3*, *MAD1*, and *MAD2* constructs were generated by previously (Komaki and Schnittger, 2017). The subcloned cDNAs were subsequently integrated into the *pGBT9-GW* (DNA-BD) or *pGAD424-GW* (AD) vectors by LR recombination reactions. The resulting constructs were transformed into the yeast strain AH109. Transformants were spotted onto control (-TL), moderately selective (-TLH), and severely selective (-TLHA) media and photographed after incubation at 30°C for 3 days. Primer pairs for plasmid construction are described in Supplemental Table S4.

### Accession Numbers

Sequence data from this article can be found under the following accession numbers: *BMF1* (At2g20635), *BMF2* (At2g33560), *BMF3* (At5g05510), *BUB3.1* (At3g19590), *BUB3.2* (At1g49910), *BUB3.3* (At1g69400), *MAD1* (At5g49880), *MAD2* (At3g25980), *MPS1* (At1g77720). *MpBMF* (Mapoly0009s0056), *MpCENP-C* (Mapoly0008s0161), *MpKNL1* (Mapoly0030s0102). The accession numbers of Arabidopsis core kinetochore genes are listed in Table 1.

## SUPPLEMENTAL DATA

The following supplemental materials are available.

Supplemental Figure S1. Y2H assay. Interaction between Arabidopsis core kinetochore components.

Supplemental Figure S2. Y2H assay. Interaction between core kinetochore and SAC components.

Supplemental Figure S3. Y2H assay. Interaction among Arabidopsis SAC components.

Supplemental Figure S4. Y2H assay. Interaction between KNL1 and its interactors.

Supplemental Figure S5. The structural prediction of KNL1 using AlphaFold2 and 3.

Supplemental Figure S6. Sequence alignment of N-terminus of CENP-C.

Supplemental Table S1. Immunoprecipitated protein list.

Supplemental Table S2. Result of GO enrichment analysis.

Supplemental Table S3. Identified phosphopeptides of kinetochore proteins.

Supplemental Table S4. Primers used in this study.

Supplemental Movie S1. Subcellular localization of CENP-S:GFP during mitosis.

Supplemental Movie S2. Subcellular localization of CENP-X:GFP during mitosis.

Supplemental Movie S3. Subcellular localization of CENP-O:GFP during mitosis.

Supplemental Movie S4. Subcellular localization of GFP: CENP-C during mitosis.

Supplemental Movie S5. Subcellular localization of GFP:NDC80 during mitosis.

Supplemental Movie S6. Subcellular localization of NUF2:GFP during mitosis.

Supplemental Movie S7. Subcellular localization of SPC24.1:GFP during mitosis.

Supplemental Movie S8. Subcellular localization of SPC24.2:GFP during mitosis.

Supplemental Movie S9. Subcellular localization of SPC25:GFP during mitosis.

Supplemental Movie S10. Subcellular localization of MIS12:GFP during mitosis.

Supplemental Movie S11. Subcellular localization of DSN1:GFP during mitosis.

Supplemental Movie S12. Subcellular localization of NNF1:GFP during mitosis.

Supplemental Movie S13. Subcellular localization of NSL1.1:GFP during mitosis.

Supplemental Movie S14. Subcellular localization of KNL1:GFP during mitosis.

Supplemental Movie S15. Subcellular localization of ZWINT1.1:GFP during mitosis.

## Acknowledgments and Funding

We thank Arp Schnittger for critical reading and comments on the manuscript, and Bo Liu and Xingguang Deng for helpful discussions. This work was supported by the Japan Society for the Promotion of Science (KAKENHI grant no. 24K09503) to S.K., the Hungarian Research Funding (NKFI-139202) to Z.M.

## Author Contributions

A.P.-S. performed immunoprecipitation assay and analyzed the data. Z.M. analyzed and discussed the data, and revised and edited the manuscript. S.K. conceived the project, conducted most of the experiments, and wrote the manuscript.

